# Likelihood-Based Inference and Model Selection for Stochastic Gene Expression in Probability-Generating-Function Space

**DOI:** 10.64898/2026.08.24.746673

**Authors:** Yiling Wang, Max Tomlinson, Zhanpeng Shu, Kim B. McAuley, Edward Z. Cao

**Affiliations:** State Key Laboratory of Bioreactor Engineering, East China University of Science and Technology, China; Department of Chemical Engineering, Queen’s University, Canada; Department of Chemical and Biological Engineering, Hong Kong University of Science and Technology, Hong Kong

## Abstract

Selecting stochastic gene-expression models from single-cell counts requires accurate parameter inference and efficient model selection. Likelihood methods in count space can be costly when full stationary count distributions are unavailable, whereas approximate methods may lose accuracy. Probability generating functions (PGFs) offer a compact analytical alternative, but existing PGF workflows are generally not likelihood based and therefore rely on computationally intensive cross-validation. We develop a likelihood-based PGF framework for both tasks. Correlated empirical PGF values are used to construct a Gaussian quasi-likelihood for parameter inference and PGF-based Bayesian information criterion (BIC) for model selection. We show that the empirical PGF is exactly unbiased and that the parameter estimator is consistent, converges at the inverse-square-root sample-size rate, and is first-order asymptotically unbiased. For large samples and a uniquely preferred model, PGF-BIC selects the same model as leave-one-out cross-validation in PGF space.

**Relevance to Life Sciences:** Cell-to-cell variability in mRNA abundance contains information about the stochastic mechanisms regulating gene expression. Distinguishing constitutive production from transcriptional bursting and promoter switching is therefore important for interpreting single-cell RNA measurements, but conventional likelihood calculations and repeated cross-validation can be computationally expensive when many genes and candidate models are considered. The proposed PGF-BIC framework provides a scalable approach for parameter inference and model selection directly from single-cell count data. Its application to MERFISH nuclear and cytoplasmic mRNA counts enables gene-wise comparison of delayed Poisson and delayed Telegraph models and produces classifications that can be compared with those obtained by tenfold PGF cross-validation. The method supports efficient screening of stochastic gene-expression models while recognizing that selection of a statistical model does not by itself establish a unique molecular mechanism.

**Mathematical Content:** The empirical probability generating function (PGF), evaluated at a fixed set of collocation points, is represented as one correlated *m*-dimensional summary vector. A multivariate central limit theorem motivates a covariance-aware Gaussian quasi-likelihood whose maximizer is the weighted minimum-distance estimator used throughout the paper. Exact unbiasedness of the empirical PGF is proved, together with consistency, root-*n* asymptotic normality, and first-order asymptotic unbiasedness of the parameter estimator under the stated identification, smoothness, and uniform-integrability conditions. A Laplace approximation to the integrated quasi-likelihood yields PGF-BIC, comprising the fitted quadratic discrepancy and the penalty *p* log *n*. For a finite candidate set, regularity and positive separation in both count and fixed-grid PGF spaces ensure that PGF-BIC and count-space BIC asymptotically select the same smallest correct candidate. Leave-one-out (LOO) cross-validation is also formulated entirely in PGF space. With a uniquely separated population minimizer, PGF-LOO and PGF-BIC select the same candidate with probability tending to one.

## 1. Introduction

Single-cell RNA counts provide information about how genes are regulated. Constitutive production, effective bursting, and explicit switching of a promoter between inactive and active states can produce different count patterns [14, 22, 23, 30]. These patterns often overlap, however, especially when other differences between cells are present [5, 21, 31]. Analyzing such data therefore involves two linked tasks. First, the parameters of each candidate model must be inferred from the observed counts. Second, the fitted models must be compared to determine which one describes the data adequately without unnecessary complexity.

Likelihood-based methods provide a natural solution when the full stationary count distribution is available. For many stochastic reaction models, however, obtaining this distribution requires solving or approximating the chemical master equation or repeatedly running stochastic simulations [9, 17, 19]. These calculations can become expensive when several models must be fitted to thousands of genes. Cheaper approaches also have limitations. Moment methods may discard differences in distributional shape, while likelihood-free and synthetic-likelihood methods depend on simulation budgets, selected summaries, and tuning choices [7, 16, 25, 33, 40]. The central challenge is therefore to retain information about the count distribution while keeping parameter inference and model selection computationally manageable.

The probability generating function (PGF) is a promising way to meet this challenge. A PGF represents the complete count distribution through a single function and can sometimes be evaluated analytically even when a large table of count probabilities is difficult to construct. Its empirical counterpart is obtained directly by averaging simple quantities over the observed cells. PGF-based estimators have a long history in count-data analysis [18, 13, 32], and recent work has shown that analytical PGFs can accelerate inference for stochastic biochemical models [15, 36].

Recent biochemical PGF workflows usually infer parameters by minimizing a distance between empirical and theoretical PGF values rather than by maximizing a sampling likelihood. As a result, model assessment commonly relies on cross-validation [15, 36]. Cross-validation is useful but requires every candidate model to be refitted on several subsets of the data. This repeated fitting amplifies the computational cost, particularly in genome-scale applications. A further statistical difficulty is that PGF values evaluated at different points are calculated from the same cells. They therefore move together and must be treated as correlated components of one data summary rather than as independent observations.

Here we develop a likelihood-based PGF framework that addresses both issues. We treat the empirical PGF evaluated on a fixed grid as one correlated random vector and use its sampling covariance to construct a Gaussian quasi-likelihood. The Gaussian approximation describes the empirical PGF, not the original count distribution. For parameter inference, optimizing this quasi-likelihood gives the covariance-weighted estimator in Eq. (2.4), which is the main estimator used throughout the paper. For model selection, the same quasi-likelihood yields a PGF version of the Bayesian information criterion, which we call PGF-BIC. Thus, parameter inference and model selection are carried out within one framework, and each candidate needs to be fitted only once to the full data.

We establish the statistical properties of both steps. The empirical PGF is exactly unbiased. Under identification and smoothness conditions, the parameter estimator is consistent, its error decreases in proportion to the inverse square root of the number of cells, and it is first-order asymptotically unbiased under an additional uniform-integrability condition. We then show that, under regularity and finite-grid separation, PGF-BIC and count-space BIC select the same smallest correct candidate with probability approaching one. We also compare PGF-BIC with leave-one-out cross-validation (LOO) performed in the same PGF space. When one candidate has a uniquely better limiting PGF fit, PGF-BIC and PGF-space LOO select the same model as the sample size becomes large. Finally, we apply PGF-BIC to multiplexed error-robust fluorescence in situ hybridization (MERFISH) data and find close agreement between its gene-level model assignments and those obtained using tenfold PGF cross-validation [4, 41, 36].

## 2. From cell counts to the PGF-based parameter estimator

### 2.1. The univariate count setting

The construction of the PGF-based parameter estimator comprises three components: a model PGF, an empirical PGF obtained by averaging over cells, and a fixed collocation map that transforms both functions into comparable vectors. Let *X*_1_, …, *X*_*n*_ denote the count observations from *n* independent cells, with each *X*_*i*_ drawn from a distribution *P*_0_ supported on the nonnegative integers. Throughout the paper, *n* denotes the number of independent cells, equivalently the sample size. For parametric inference, assume that 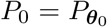 for some interior point ***θ***_0_ ∈ Θ ⊂ ℝ ^*p*^, and that the model PGF *G*_***θ***_(*z*) can be evaluated without reconstructing the entire probability mass function. For a nonnegative integer-valued random variable, the PGF is defined as

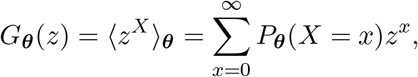

for *z* ∈ [0, 1]. Throughout, a subscript 0 denotes a quantity associated with the data-generating distribution, and ⟨·⟩_0_ denotes expectation under *P*_0_. Rather than evaluating the PGF at every *z*, we use a finite grid of evaluation points, referred to as collocation points. The number of collocation points is denoted as *m*. The grid is selected independently of the data and is shared by all candidate models. In all fixed-grid asymptotic statements, the number of cells satisfies *n* → ∞, whereas *m* and the collocation points *z*_1_, …, *z*_*m*_ remain fixed:

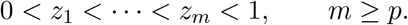

The condition *m* ≥ *p* is necessary for the full-column-rank requirement introduced below, although it is not sufficient for identification. In model-selection settings, *m* must be at least as large as the maximum parameter dimension among the candidate models. The endpoint *z* = 1 is excluded because 1^*X*^ = 1 almost surely and therefore has zero variance.

At any fixed *z*, the data provide the empirical counterpart

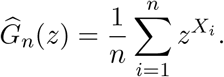

Evaluating these quantities at the collocation points yields a feature vector for each cell, together with the corresponding model and empirical vectors

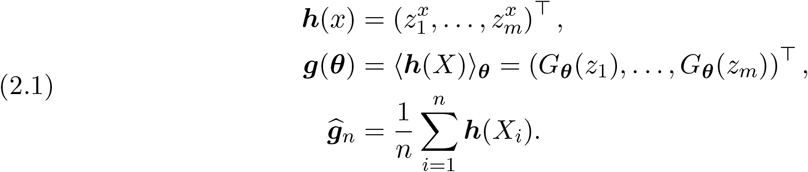

To streamline notation throughout, let ***g***_0_ = ⟨***h***(*X*)⟩_0_ denote the true PGF vector and 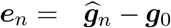 its empirical estimation error. Under correct model specification, ***g***_0_ = ***g***(***θ***_0_). Because all *m* coordinates are computed from the same cells, they constitute a single correlated vector rather than *m* independent observations. The covariance matrix of a cell-level feature vector is

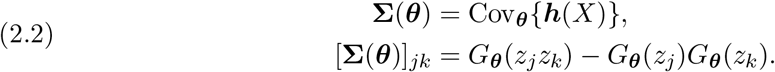

To see the second identity in Eq. (2.2), note that the *j*th and *k*th coordinates of the feature vector from the same cell are 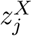 and 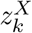. Therefore,

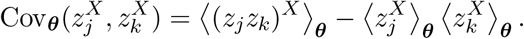

Because 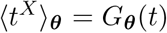, this expression gives the second line of Eq. (2.2). The off-diagonal entries are generally nonzero because all PGF coordinates are evaluated using the same cell-level count. In the GMM interpretation, **Σ**(***θ***) is the covariance matrix of the moment vector and determines how the correlated PGF coordinates should be weighted [10]. Under the data-generating distribution, let **Σ**_0_ = Cov_0_{***h***(*X*)}. It follows that the covariance matrix of 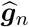 is **Σ**_0_*/n*, rather than **Σ**_0_. The corresponding empirical estimator of the single-cell covariance matrix is

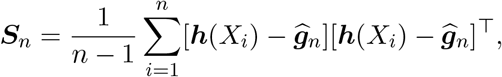

which provides a model-agnostic estimate of **Σ**_0_. Because every coordinate of ***h***(*X*) is bounded, the multivariate central limit theorem yields

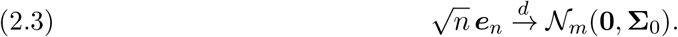

The unregularized inverse 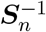 exists only if *n > m* and the retained cell-level features have full column rank.

Equation (2.3) motivates a working likelihood for the empirical PGF vector, but it does not imply that the collocation coordinates are independent observations. Let ***W***_*n*_ be a positive-definite weight matrix that is common to all parameter values and, in model-selection settings, to all candidate models. By default, we set 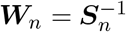. When ***S***_*n*_ is ill-conditioned, we apply ridge regularization by setting ***W***_*n*_ = (***S***_*n*_ + *λ***I**_*m*_)^−1^, where *λ >* 0 and **I**_*m*_ denotes the *m* × *m* identity matrix [37]. The resulting plug-in working density is

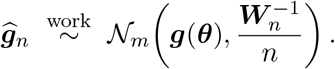

This is a Gaussian quasi-likelihood for 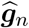, not an exact finite-sample likelihood for the counts,

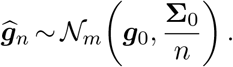

It resembles synthetic likelihood for summary statistics, but here the PGF mean is evaluated analytically and the covariance is estimated from bounded cell-level quantities rather than by repeated simulation [40]. Up to a term *c*_*n*_ that does not depend on ***θ***, the quasi-log-likelihood is

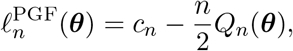

with

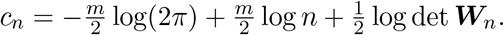

The weighted PGF discrepancy is

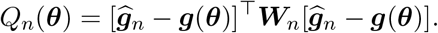

Maximizing the working likelihood is therefore equivalent to minimizing the corresponding weighted distance. The estimator used throughout the paper is thus

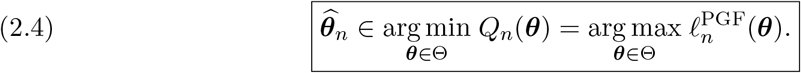

As Eq. (2.4) makes clear, the estimator is governed by two objects: the model surface ***g***(Θ) and the weighting matrix ***W***_*n*_. The former determines whether the parameters are identifiable from the chosen collocation grid, whereas the latter controls how discrepancies along correlated PGF directions are weighted. Multiplying the objective function by *n* does not change its minimizer, but this factor is essential to the curvature of the quasi-likelihood and to the model-selection penalty derived later. Equation (2.4) is an empirical-PGF minimum-distance, or generalized method of moments (GMM), estimator [10, 13, 15, 18, 32]. Under correct specification, the default choice 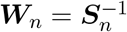 is first-order efficient among estimators based on the chosen PGF moments. The workflow of the PGF-based parameter inference method is summarized in Fig. 1. All subsequent PGF fits in the paper use this estimator.

**Figure 1:**
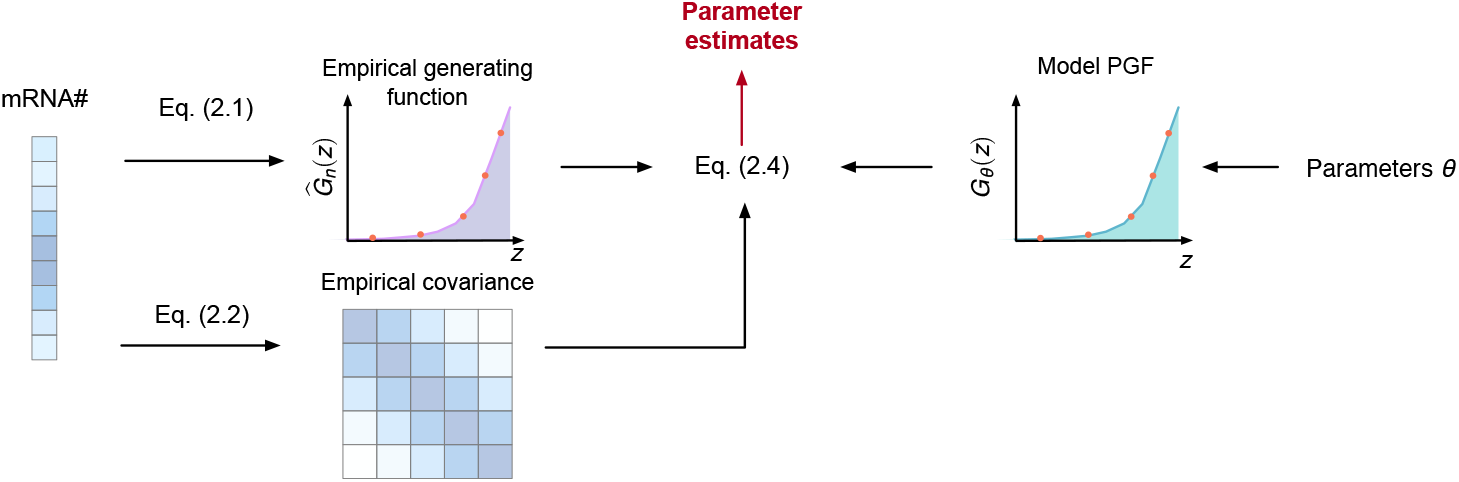
Schematic of covariance-aware parameter inference in PGF space. Single-cell mRNA counts are used to calculate the empirical PGF at a fixed set of collocation points (Eq. (2.1)) and the covariance among these empirical values (Eq. (2.2)). For a trial parameter vector ***θ***, the model PGF is evaluated at the same points. Equation (2.4) compares the empirical and model PGF vectors while accounting for their covariance, and minimizing the resulting weighted discrepancy yields the parameter estimate. The red points denote the collocation points used in the comparison.

### 2.2. Extension to multivariate count distributions

Recent advances in single-cell transcriptomics enable the simultaneous quantification of biologically distinct RNA populations within the same cell. MERFISH and related multiplexed imaging approaches can resolve spatially distinct RNA pools and, with intron-targeting probe designs, nascent transcripts, whereas metabolic RNA-labeling assays distinguish newly synthesized from pre-existing RNA [2, 6, 12, 29, 41]. Motivated by these data, we extend our method to a joint-count formulation. For a multivariate count For cell *i*, let ***X***_*i*_ = (*X*_*i*1_, …, *X*_*iv*_)^T^ be a *v*-dimensional count vector, where *v* is the number of jointly modeled count components (for example, molecular species or cellular compartments). The corresponding joint model PGF is

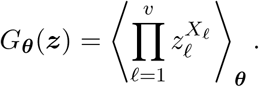

Choose collocation points ***z***_*j*_ ∈ (0, 1)^*v*^ and define the corresponding bounded joint features by

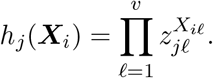

When the joint model PGF is used, the estimator in Eq. (2.4) retains the same form, with covariance entries given by

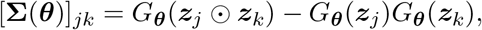

where ⊙ denotes componentwise multiplication. Thus, the joint-count formulation introduces no new estimator; only the bounded cell-level feature map and its model-implied expectation change. For notational simplicity, the remainder of the theoretical development uses the univariate-count formulation, with *n* always denoting the number of independent cells.

## 3. Why the PGF estimator works

Equation (2.4) first estimates the PGF by averaging over cells and then maps that estimate back to model parameters. The first step is simple because every component of ***h***(*X*) is bounded: the empirical PGF is exactly unbiased and has sampling error of order *n*^−1*/*2^. The second step requires the chosen grid to identify the true parameter. A full-rank Jacobian then transfers the same sampling rate to the parameter estimate. Because this mapping is nonlinear, exact finite-sample unbiasedness belongs to 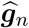, not generally to 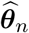. The parameter estimator is instead first-order asymptotically unbiased under the additional condition stated below. The theorem makes these claims precise using standard minimum-distance and GMM arguments [10, 20].

Theorem 3.1 (Unbiased empirical PGF and root-*n* inference). *Let* 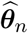 *be any measurable global minimizer in Eq*. (2.4). *Suppose that:*

1. Θ *is compact*, ***θ***_0_ *is an interior point of* Θ, *and X*_1_, …, *X*_*n*_ *are independent and identically distributed (i*.*i*.*d*.*) according to* 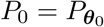;
2. ***g*** *is continuous on* Θ, *and* ***g***(***θ***) = ***g***_0_ *implies* ***θ*** = ***θ***_0_;
3. ***g*** *is continuously differentiable in a neighborhood of* ***θ***_0_, *and the m* × *p Jacobian*

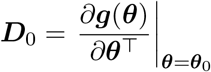

*has full column rank;*
4. 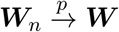, *where* ***W*** *is positive definite*.
*Then the following statements hold*.
  i. *The empirical PGF vector is exactly unbiased, with*

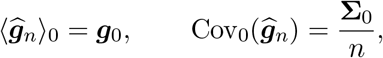

*and satisfies Eq*. (2.3).
  ii. *The parameter estimator is consistent and admits the expansion*

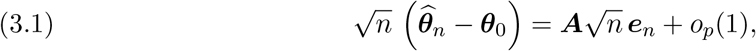

*where*

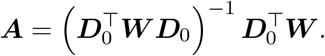

*Consequently*,

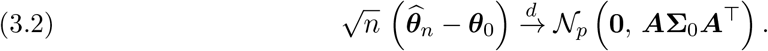

*If* **Σ**_0_ *is nonsingular, the choice* 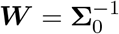*reduces the limiting covariance to*

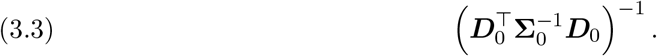
  iii. *If, in addition, the scalar sequence* 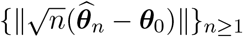 *is uniformly integrable, then the parameter estimator is first-order asymptotically unbiased*,

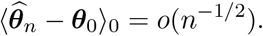
  *Proof*. The proof follows the same data-to-parameter path as the construction: first the empirical PGF, then the minimum of its quadratic criterion, and finally the local inversion that produces the parameter estimate. For coordinate *j*, the definition of the empirical PGF and linearity of expectation give

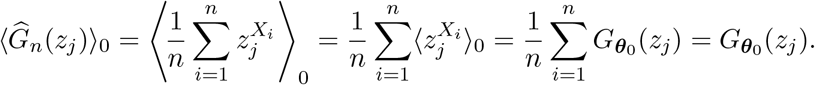

Stacking these identities for *j* = 1, …, *m* gives 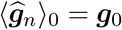. This equality is exact for every *n*; no normal approximation has been used.

Independence of the cells removes all cross-cell covariance terms. Hence

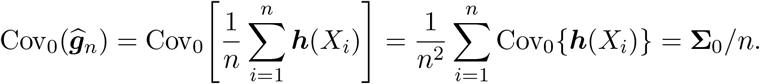

The (*j, k*) entry of the one-cell covariance is

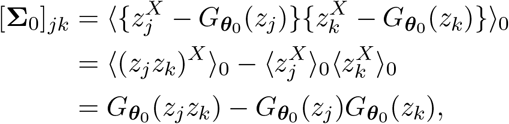

which agrees with Eq. (2.2).

Because 0 *< z*_*j*_ *<* 1 and *X* is nonnegative, 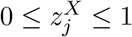. Thus ***h***(*X*_*i*_) is a bounded random vector. To make the multivariate central limit theorem explicit, fix ***a*** ∈ ℝ^*m*^. The variables ***a***^⊤^[***h***(*X*_*i*_) − ***g***_0_] for *i* = 1, …, *n*, are i.i.d., centered, and bounded. The scalar central limit theorem gives

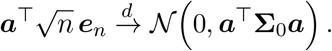

It follows by the Cramér–Wold device [34, p. 16] that

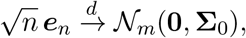

because every linear combination of the PGF errors has the corresponding Gaussian limit. This proves part (i). The covariance **Σ**_0_ need not be nonsingular here; the limiting normal law may be supported on a lower-dimensional subspace.

The law of large numbers gives 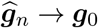 almost surely, and hence ***e***_*n*_ = *o*_*p*_(1). Define

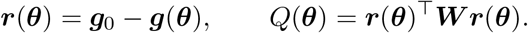

Every coordinate of both ***g***_0_ and ***g***(***θ***) lies in [0, 1]. Therefore, 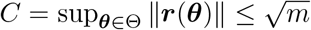. Since 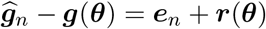, direct expansion gives

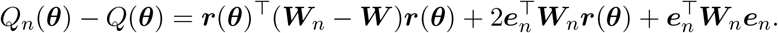

Using |***u***^⊤^***Mv***| ≤ ∥***M*** ∥ ∥***u***∥ ∥***v***∥, we obtain the uniform bound

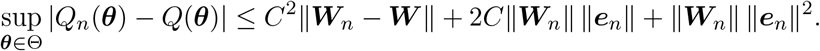

Assumption 4 implies ∥***W***_*n*_ − ***W*** ∥ = *o*_*p*_(1) and ∥***W***_*n*_∥ = *O*_*p*_(1). Together with ***e***_*n*_ = *o*_*p*_(1), the bound proves

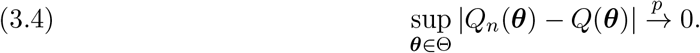

For each sample, *Q*_*n*_ is continuous on compact Θ, so its global argmin is nonempty. At the population level, positive definiteness of ***W*** gives

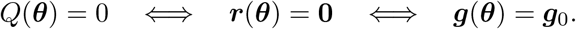

By identification, ***θ***_0_ is therefore the unique minimizer of *Q*, and *Q*(***θ***_0_) = 0.

It remains to show that a minimizer of the random function *Q*_*n*_ must be close to this unique population minimizer. Fix *ε >* 0 and collect all parameter values that are not within *ε* of the truth in

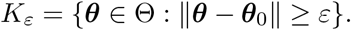

If *K*_*ε*_ is empty, the desired probability is already zero. If it is nonempty, then it is a closed subset of compact Θ, and hence is compact. Continuity of *Q* means that its infimum over *K*_*ε*_ is attained at some ***θ***_*ε*_ ∈ *K*_*ε*_. Because ***θ***_0_ ∈*/ K*_*ε*_ and ***θ***_0_ is the unique global minimizer,

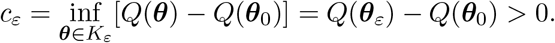

Thus *c*_*ε*_ is the positive population-criterion gap that separates the truth from every parameter at least *ε* away.

Now set δ_*n*_ = sup***θ***∈Θ |*Q*_*n*_(***θ***) − *Q*(***θ***)|. By its definition, simultaneously for every ***θ*** ∈ Θ,

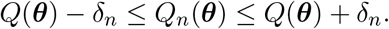

At the true parameter, this bound gives

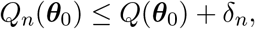

while the corresponding lower bound over *K*_*ε*_ is

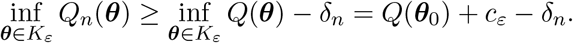

Subtracting these two bounds gives

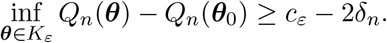

If δ_*n*_ *< c*_*ε*_*/*2, the uniform sampling error is too small to close the population gap: *c*_*ε*_ − 2δ_*n*_ *>* 0. It follows that every parameter at least *ε* away from ***θ***_0_ has a larger sample criterion than ***θ***_0_.

On the other hand, because 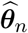 is a global minimizer,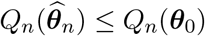. Hence 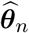 cannot be one of the parameter values that are at least *ε* away; it must satisfy 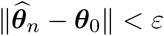. Thus, the estimator can lie outside this neighborhood only when δ_*n*_ ≥ *c*_*ε*_*/*2:

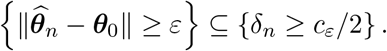

Consequently,

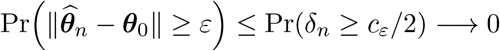

because 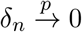 by Eq. (3.4). This proves 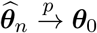, equivalently the basic extremum-estimator consistency result of Ref. [20, Theorem 2.1, p. 2121]. In words, uniform convergence places the entire sample criterion inside a shrinking band around the population criterion; once that band is narrower than half of the separation gap, its minimizer cannot lie far from ***θ***_0_.

Choose an open ball around ***θ***_0_ whose closure lies both in the interior of Θ and in the neighborhood where ***g*** is continuously differentiable. Consistency places 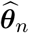 in this ball with probability approaching one. On that event it is an interior minimizer, so differentiating *Q*_*n*_ gives

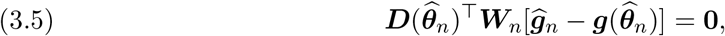

where ***D***(***θ***) = ∂***g***(***θ***)*/*∂***θ***^T^. The omitted factor −2 does not affect the first-order condition.

Set 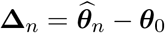 and define the Jacobian averaged along the line segment between the true and estimated parameters:

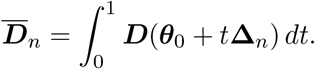

Applying the fundamental theorem of calculus coordinate by coordinate gives the exact identity

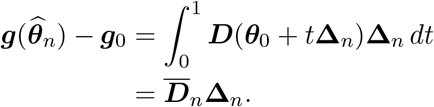

It follows that

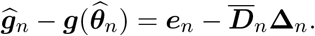

Substitution into Eq. (3.5) yields

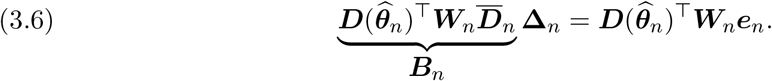

This equality shows how the empirical-PGF error ***e***_*n*_ is transferred to the parameter error **Δ**_*n*_ through a locally invertible matrix.

Consistency and continuity of ***D*** imply

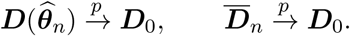

Together with 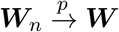, these limits give

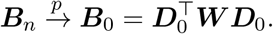

For every nonzero ***v*** ∈ ℝ^*p*^,

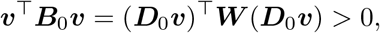

because ***D***_0_ has full column rank and ***W*** is positive definite. Thus ***B***_0_ is positive definite. Continuity of matrix inversion then implies that ***B***_*n*_ is invertible with probability approaching one and 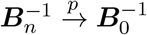.

Multiplying Eq. (3.6) by 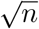 and solving for 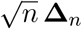 gives

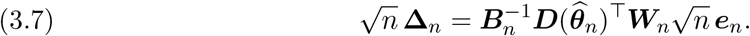

Write the matrix multiplying 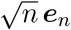 as

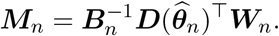

It converges in probability to

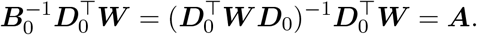

Thus ***M***_*n*_ − ***A*** = *o*_*p*_(1). Step 1 also implies 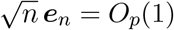, so

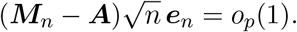

Replacing the random matrix in Eq. (3.7) by ***A*** therefore gives

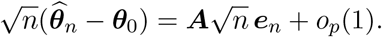

This proves Eq. (3.1). The claim (i) and Slutsky’s lemma [34, Lemma 2.8, p. 11] then give Eq. (3.2).

For 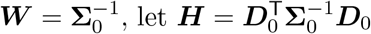. Direct substitution shows

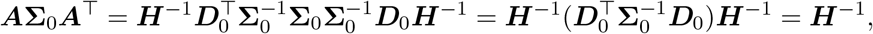

which is Eq. (3.3).

The final claim concerns bias at the root-*n* scale; it does not assert finite-sample unbiasedness of the parameter estimator. Let

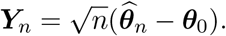

Part (ii) gives 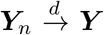, where 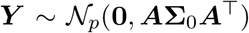. By assumption, {∥***Y***_*n*_∥} is uniformly integrable. Since |*Y*_*n,k*_| ≤ ∥***Y***_*n*_∥ for every coordinate *k*, each scalar sequence {*Y*_*n,k*_} is also uniformly integrable. Convergence in distribution together with uniform integrability therefore gives convergence of first moments [34]:

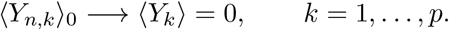

Thus ⟨***Y***_*n*_⟩_0_ → **0**, and dividing by 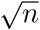 gives

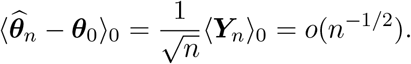

This proves part (iii).

Theorem 3.1 establishes the large-sample properties of the PGF estimator. We next examine whether its predicted *n*^−1*/*2^ convergence rate is visible at finite sample sizes. Figure 2 presents results for a transcriptional bursty model [24] and the telegraph model [3]. Across all six model–parameter settings, the mean Euclidean estimation error decreases approximately linearly on the log–log scale and closely follows the reference slope of −1*/*2. These results provide numerical support for the convergence rate established in the theorem.

**Figure 2:**
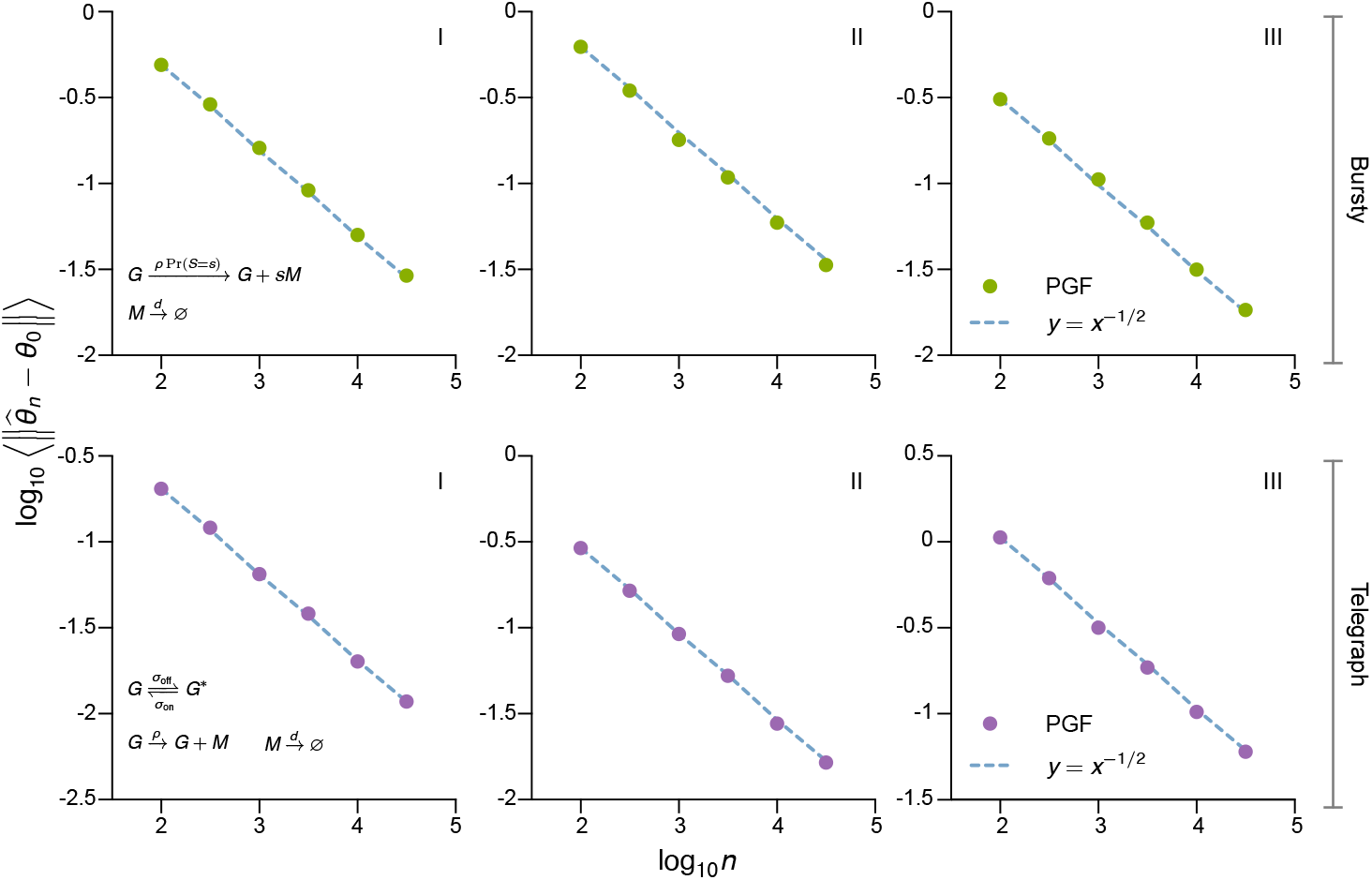
Finite-sample convergence of the PGF-based parameter estimator. Insets show the corresponding reaction schemes. The upper row considers the bursty model, in which bursts occur at rate *ρ* and have geometrically distributed size *S*, with Pr(*S* = *s*) = *b*^*s*^*/*(1 + *b*)^*s*+1^ and mean *b*. The lower row considers the telegraph model, in which the promoter switches between inactive and active states at rates *σ*_on_ and *σ*_off_, and only the active state transcribes at rate *ρ*. Transcripts degrade at rate *d* = 1 in both models. Columns I–III use (*ρ, b*) = (4, 0.4), (6, 0.6), (3, 0.5) for the bursty model and (*ρ, σ*_on_, *σ*_off_) = (8, 0.5, 0.1), (13, 1, 0.4), (25, 0.5, 1.4) for the telegraph model. Each dot shows the mean Euclidean parameter-estimation error across 1000 independent data sets simulated using `DelaySSAToolkit.jl` [8]. The empirical PGF and its covariance were evaluated at ***z*** = (0.90, 0.93, 0.96, 0.99). Dashed blue lines have slope − 1*/*2, representing the *n*^−1*/*2^ rate predicted by Theorem 3.1.

## 4. Deriving BIC in the PGF space

Equation (2.4) answers the parameter-inference question for one model: it finds the parameter value whose PGF is closest to the empirical PGF. Model selection asks a further question. If a model with more parameters fits better, is the improvement large enough to justify the extra complexity? Fit alone cannot answer this question, because additional parameters usually make it easier to follow sampling noise. We therefore use the same PGF quasi-likelihood to derive a PGF version of BIC named PGF-BIC.

No new parameter estimator is introduced.

Consider a finite collection of candidates ℳ_1_, …, ℳ_*R*_. Candidate *r* has *p*_*r*_ free parameters collected in 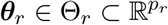. Its values on the common PGF grid form the vector

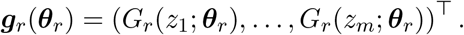

Every candidate is compared with the same empirical PGF 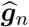, the same grid, and the same weight ***W***_*n*_. This common representation puts all fitted discrepancies on the same scale. Only the model PGF ***g***_*r*_ and the number of parameters *p*_*r*_ change between candidates. For candidate *r*, the weighted PGF discrepancy is

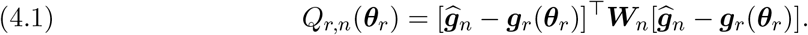

Let 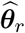 denote the parameter estimate obtained by globally minimizing *Q*_*r,n*_. This is exactly Eq. (2.4) applied to candidate *r*. We omit the sample-size subscript from the fitted parameter; hats denote PGF fits, tildes denote the count-space fits introduced later, and *n* continues to distinguish sample criteria from population criteria.

The Gaussian approximation for the empirical PGF gives candidate *r* the quasi-log-likelihood

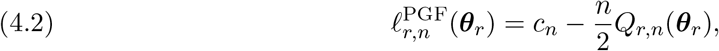

where *c*_*n*_ is a normalizing term common to all candidates. It is common because they use the same grid dimension and the same parameter-independent weight ***W***_*n*_. Minimizing *Q*_*r,n*_ is therefore the same as maximizing this quasi-likelihood. The word “quasi” is important: this is an approximate likelihood for the empirical PGF vector, not the exact likelihood of the original cell counts. The Gaussian approximation is justified locally for models that reproduce the true PGF. For a model separated from the true PGF, *Q*_*r,n*_ remains a useful weighted measure of PGF mismatch.

To describe what remains after sampling noise is removed, replace the empirical PGF and weight by their population limits. Reuse the true PGF vector ***g***_0_ from Section 2, let 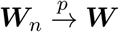, and define ***d***_*r*_(***θ***_*r*_) = ***g***_0_ − ***g***_*r*_(***θ***_*r*_). The population discrepancy is

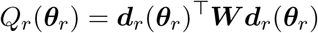

and is the large-sample version of *Q*_*r,n*_. Let 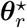 be its unique minimizer. The ordinary BIC penalty arises in the regular local regime: this minimizer must be interior and isolated, and the negative Hessian of the quasi-log-likelihood must grow linearly with the number of cells. The precise curvature condition is stated below. Accordingly, suppose that 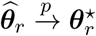, that the quasi-likelihood has a uniform local quadratic expansion on *n*^−1*/*2^-scale neighborhoods, and that its integrated mass outside a fixed neighborhood of 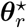 is negligible.

BIC is obtained by integrating over parameter uncertainty instead of looking only at the best-fitting value. For the derivation, assign candidate *r* a proper prior density *π*_*r*_(***θ***_*r*_) that is continuous and positive near 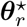. This density supplies a smooth weight over possible parameter values. The resulting integrated Gaussian quasi-likelihood is

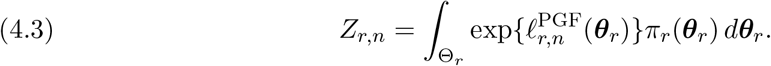

The width of this region is controlled by the local curvature: how quickly the quasi-log-likelihood decreases as the parameter moves away from its fitted value. Define the curvature matrix, also called the observed quasi-information, by

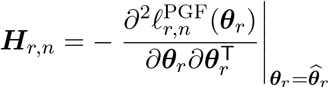

and assume the required smoothness and positive local curvature:

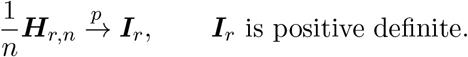

At a smooth minimum, 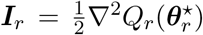. For a correctly specified candidate, this is the candidate-specific version of 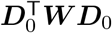 in Theorem 3.1. Positive definiteness of ***I***_*r*_ is imposed rather than inferred, because it can fail at a boundary or singular point.

The determinant of a matrix is the product of its eigenvalues. Because ***H***_*r,n*_ has *p*_*r*_ parameter directions and each eigenvalue grows in proportion to *n*, its determinant grows in proportion to 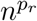. Equivalently,

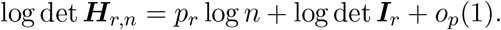

This *p*_*r*_ log *n* term becomes the complexity penalty.

A Laplace approximation evaluates the integral in Eq. (4.3) by replacing the quasi-likelihood near its peak with a Gaussian shape having the same height and curvature. It gives

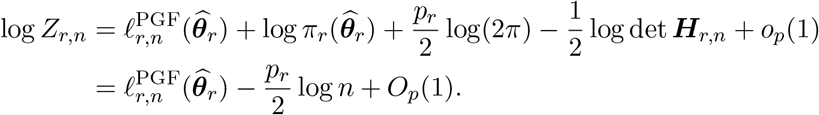

To see the final score directly, multiply the last expression by −2 and substitute Eq. (4.2):

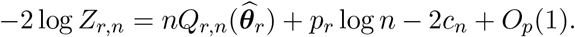

The normalization *c*_*n*_ is common to all candidates. The remaining *O*_*p*_(1) terms, which include prior and finite-curvature contributions, may differ between candidates but remain bounded in probability. Removing the common normalization and omitting these order-one terms, as in the standard BIC derivation [26], gives the implementable score

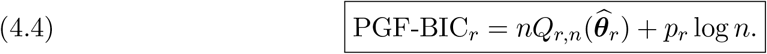

The two terms have simple roles. The first measures lack of fit in PGF space; smaller is better. The second penalizes the *p*_*r*_ fitted parameters, so a more complex model must improve the PGF fit enough to overcome its larger penalty. Because the fitted discrepancy comes directly from Eq. (2.4), parameter inference and model selection use the same full-data fit and the same covariance weight.

The selected model is the candidate with the smallest PGF-BIC:

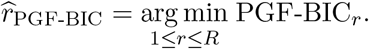

The penalty in Eq. (4.4) counts free kinetic and nuisance parameters, not collocation points. Here *n* is the number of independent cells, *p*_*r*_ is the number of independently identifiable parameters, and *m* is only the length of the common PGF summary. The collocation points are correlated summaries of the same cells; they are neither extra observations nor fitted parameters. Likewise, estimating one common ***W***_*n*_ from the cell-level features does not add model-specific parameters. The criterion in Eq. (4.4) only uses the leading *n*-order fit and *p*_*r*_ log *n* curvature terms; order-one terms can still matter in small samples or nearly tied comparisons.

## 5. Selection equivalence with count-space BIC

Having derived the score Eq. (4.4), we next ask whether it and count-space BIC choose the same model label, not whether their numerical values are equal.

The derivation above does not by itself guarantee agreement with count-space BIC. Agreement of selected labels requires two kinds of separation. A false candidate must remain detectably false in both the full count law and the fixed-grid PGF summary; among the correct candidates, the shared dimension penalty must then single out one smallest model. We now state conditions under which these two steps succeed. Let *f*_0_ be the true count mass function and *f*_*r*_(*x*; ***θ***_*r*_) the mass function under ℳ_*r*_. The sample log-likelihood, its count-space maximum-likelihood estimator (MLE), and the ordinary BIC are

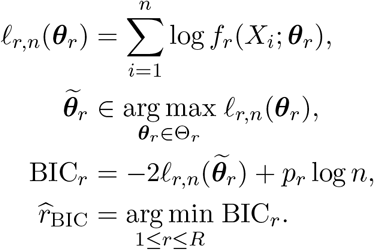

A hat continues to denote a PGF fit, while a tilde denotes the count-space MLE. All fits are measurable global optima, and a fixed rule resolves exact ties.

The two required separations are measured by parallel population discrepancies. Reusing the log-likelihood symbol without the sample-size index, set

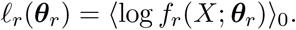

The count-space Kullback–Leibler (KL) discrepancy is

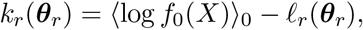

and its PGF-space counterpart is the previously defined *Q*_*r*_. Their minimum values are

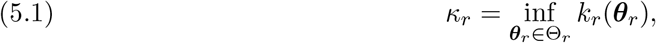

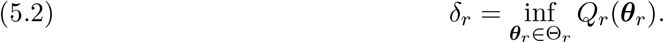

Also let

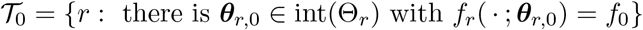

be the set of correctly specified candidates. This construction combines the regular BIC logic of Schwarz and Haughton [11, 26], the KL analysis of misspecified likelihood [27, 38], and the finite-summary separation used in GMM model selection [1].

The assumptions below play three roles: they make the optimized sample criteria converge uniformly to their population versions, keep every false candidate a positive distance from the truth in both representations, and retain regular likelihood and PGF curvature for the correct candidates.

### Theorem 5.1

(Selection equivalence of PGF-BIC and count-space BIC). *Suppose that: 1. The candidate collection is finite, the observations are i*.*i*.*d. from f*_0_, *and the set* T_0_ *defined above is nonempty. There is a unique smallest correct candidate r*_0_, *meaning* 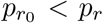 *for every r* ∈ T_0_ \ {*r*_0_}.

2. *Each parameter space* Θ_*r*_ *is compact, and the same fixed points z*_1_, …, *z*_*m*_ ∈ (0, 1) *are used for every model. Each map* ***g***_*r*_ *is continuous, and every fit* 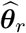 *is a measurable global instance of Eq*. (2.4), *equivalently a global minimizer of Q*_*r,n*_ *in Eq*. (4.1). *The common* ***W***_*n*_ *is symmetric positive definite and* 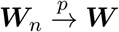 *for a fixed positive-definite matrix* ***W***. *In particular, if* 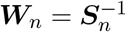, *the feature covariance* **Σ**_0_ = Cov_0_{***h***(*X*)} *is positive definite, so* 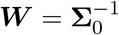.
3. *Every false candidate r* ∈*/* T_0_ *is separated in both representations:* 0 *< κ*_*r*_ *<* ∞ *and* δ_*r*_ *>* 0.
4. *We have* ⟨| log *f*_0_(*X*)|⟩_0_ *<* ∞. *For every r, the model mass function is positive for every* ***θ***_*r*_ ∈ Θ_*r*_, *almost surely under f*_0_, *and ℓ*_*r*_ *is finite on* Θ_*r*_, *with*

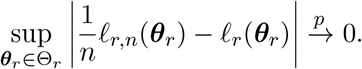

*This means that the sample log-likelihood per cell approaches its expected value simultaneously over the whole parameter space. One sufficient set of conditions* [20] *is almost-sure continuity in* ***θ***_*r*_, *together with*

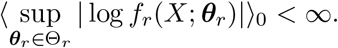

*For each correct candidate*, ***θ***_*r*,0_ *is globally identified, the global MLE is consistent for it, and the count likelihood is locally asymptotically normal with nonsingular information. In particular, assume*

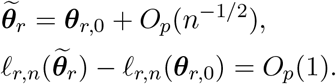

*The PGF Jacobian has full column rank at* ***θ***_*r*,0_, *as required by the Laplace derivation, and the same root-n-identified dimension p*_*r*_ *is used in both penalties*.
*Then, as n* → ∞,

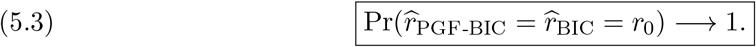

*In plain terms, the probability that both procedures select the unique smallest correct candidate approaches one. More specifically, with* 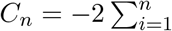 log *f*_0_(*X*_*i*_), *for each fixed r*,

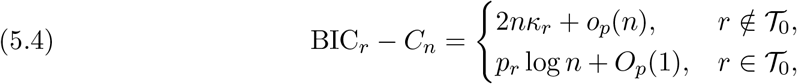

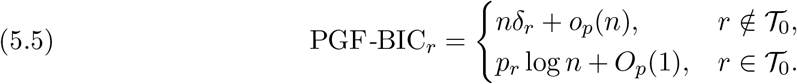

*Here C*_*n*_ *is a count-likelihood baseline shared by every candidate, so subtracting it does not change which count-space BIC is smallest. These equations summarize the central comparison: an inadequate model pays a loss proportional to n, whereas a correct model has only fixed-scale fitting variation plus the p*_*r*_ log *n penalty. The proof therefore compares three scales:*

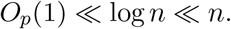

*The order-n loss removes inadequate candidates; among the remaining correct candidates, the order-*log *n penalty selects the one with the fewest parameters*.

*Proof*. The proof has three steps. We first compare inadequate and correct candidates in count space, then repeat the comparison in PGF space, and finally translate the resulting score sizes into model selection.

We begin with count space. The assumed uniform law of large numbers gives

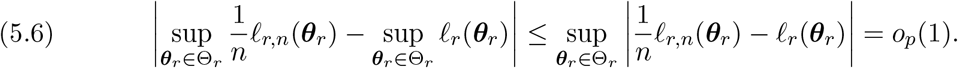

The left-hand side compares the best sample log-likelihood per cell with the best expected log-likelihood per cell.

The inequality says that maximizing cannot make this difference larger than the largest approximation error over the parameter space. Hence the optimized sample value approaches the optimized population value.

Also,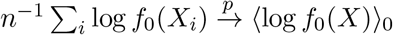. By the definition of *κ_r_*,

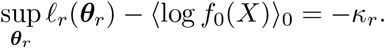

Combining Eq. (5.6) and using the fact that 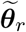 globally maximizes *ℓ*_*r,n*_ gives, for every false candidate,

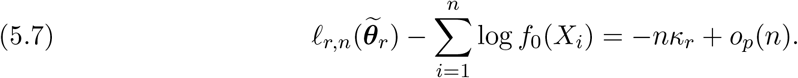

Thus a positive per-cell KL discrepancy becomes a likelihood loss proportional to *n* over the full sample. This optimized-value argument is the elementary form of the Kullback–Leibler analysis for misspecified maximum likelihood [38, Theorem 2.2, p. 4].

If *r* ∈ *T*_0_, then *ℓ*_*r,n*_(***θ***_*r*,0_) = ∑_*i*_ log *f*_0_(*X*_*i*_). Local asymptotic normality together with the stated root-*n* behavior of the regular MLE yields the *O*_*p*_(1) likelihood gain below, as in the setting of Wilks’ likelihood-ratio theorem [39, p. 62],

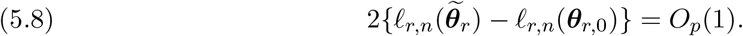

The left-hand side is the likelihood-ratio statistic comparing the fitted and true parameters; *O*_*p*_(1) means that it does not increase systematically with sample size. Adding *p*_*r*_ log *n* to Eqs. (5.7) and (5.8), and noting that *p*_*r*_ log *n* = *o*(*n*), proves Eq. (5.4). A false candidate therefore pays an order-*n* lack-of-fit cost in count space, whereas a correct candidate has only fixed-scale fitting variation plus its *p*_*r*_ log *n* penalty.

We now establish the parallel result in PGF space. Recall that 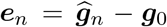 is the empirical PGF error. Because the empirical PGF is an average over *n* independent cells, ∥***e***_*n*_∥ = *O*_*p*_(*n*^−1*/*2^). For candidate *r*,

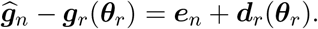

Every coordinate of ***d***_*r*_ lies in [−1, 1], so 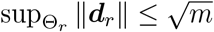. Expanding the sample quadratic and applying the standard matrix-norm bound therefore gives

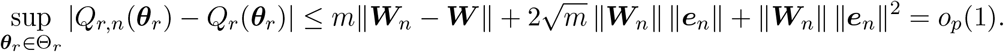

Here 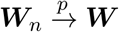 makes ∥***W***_*n*_ − ***W*** ∥ = *o*_*p*_(1) and ∥***W***_*n*_∥ = *O*_*p*_(1). Therefore, the smallest sample PGF discrepancy approaches the smallest population PGF discrepancy. If *r* ∉ T_0_, Eq. (5.2) therefore implies

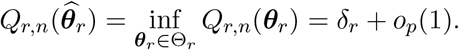

Thus the inadequate candidate retains a positive PGF mismatch even after its parameters have been optimized. Multiplication by *n*, followed by *p*_*r*_ log *n* = *o*(*n*), proves the false-model line of Eq. (5.5).

If *r* ∈ *T*_0_, then ***g***_*r*_(***θ***_*r*,0_) = ***g***_0_. At this true parameter, the PGF criterion contains only sampling error. Because 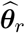 globally minimizes the criterion, its fitted value cannot exceed the value at ***θ***_*r*,0_:

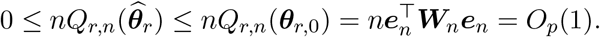

The last equality in order follows because ***e***_*n*_ = *O*_*p*_(*n*^−1*/*2^): its squared weighted norm is *O*_*p*_(*n*^−1^), and multiplication by *n* leaves a fixed-scale random term. Adding *p*_*r*_ log *n* proves the true-model line of Eq. (5.5). Notice that this step uses the proposed estimator in Eq. (2.4) directly; no auxiliary PGF estimator or model-specific covariance fit is involved.

It remains to translate these score sizes into model selection. Compare each candidate with the unique smallest correct candidate *r*_0_. For every false *r* ∉ T_0_, Eqs. (5.4) and (5.5) give

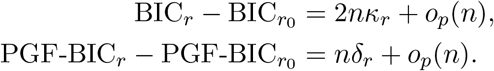

Because *κ*_*r*_ *>* 0 and δ_*r*_ *>* 0, both differences are positive with probability approaching one. In either representation, the order-*n* lack-of-fit cost eventually dominates the order-log *n* parameter penalty.

For every other correct candidate *r* ∈ T_0_ \ {*r*_0_}, the fitted values differ only by fixed-scale random terms, so the leading score difference comes from the parameter penalty:

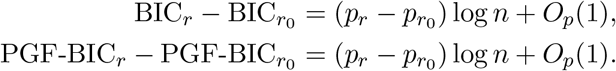

By assumption, 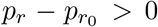. The positive 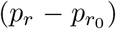 log *n* term therefore grows without bound, while the *O*_*p*_(1) fluctuation stays on a fixed scale. Both differences diverge to +∞ in probability.

We have shown that the probability that any fixed competitor beats *r*_0_ tends to zero. Because there are only finitely many candidates, the probability that at least one competitor wins is no larger than the sum of these vanishing probabilities:

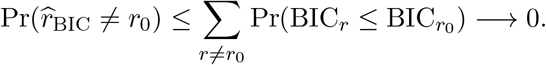

The probability of a union is no larger than the sum of the individual probabilities, so the same argument applies to PGF-BIC. The probability that both selectors equal *r*_0_ therefore approaches one, proving Eq. (5.3). The proof compares orders and signs; it does not assert equality of the two scores, their parameter estimates, or the two discrepancies *κ*_*r*_ and δ_*r*_.

### Remark 5.2

(What the equivalence does and does not say). Theorem 5.1 establishes equivalence of the selected model label, not equality of the two scores. A false model loses by 2*nκ*_*r*_ to leading order in count space and by *n*δ_*r*_ in PGF space. These two gaps need only be positive; they need not have the same numerical value. The unique-smallest-model assumption is also necessary for agreement of labels. If two correct candidates have the same smallest dimension, their *p*_*r*_ log *n* penalties are equal. Their comparison is then decided by fixed-scale random terms, and the two criteria need not choose the same label.

Theorem 5.1 establishes that count-space BIC and PGF-BIC eventually select the same model when the true model is included and every false candidate remains separated from it in both count and PGF space. The proof explains the large-sample result, but it does not show how quickly the two procedures agree when some candidates provide very similar distributions. We therefore examine the theorem numerically in a setting where the distance between competing models can be varied continuously.

The stationary telegraph distribution (Fig. 2) provides a useful example. In the classical bursting limit, it approaches a negative-binomial (NB) distribution. In a different limit, when promoter switching becomes fast relative to mRNA degradation while the active-state fraction remains fixed, it approaches a Poisson distribution. More generally, an NB distribution can closely approximate the stationary telegraph distribution over a wider intermediate parameter range, even when the system is not in the classical bursting limit [35, Secs. 2.1–2.3].

These relationships raise a natural model-selection question. If the data are generated by a telegraph model, should they be described by the full telegraph model, or can the simpler NB or Poisson model capture the observed distribution adequately? The answer depends on the sample size. With few cells, a small improvement in fit may not justify the additional parameters of the telegraph model. With more cells, differences that were previously hidden by sampling variation may become detectable.

The approximate expected Bayesian information criterion (aeBIC) was introduced to study this question without repeatedly generating and fitting many independent data sets [35, Eqs. (10)–(13) and Fig. 4]. Instead of calculating BIC for many random samples and then averaging the results, aeBIC replaces the random fitted log-likelihood by its best population value under the known generating distribution. Thus, only one population optimization is required for each candidate (Fig. 3a).

**Figure 3:**
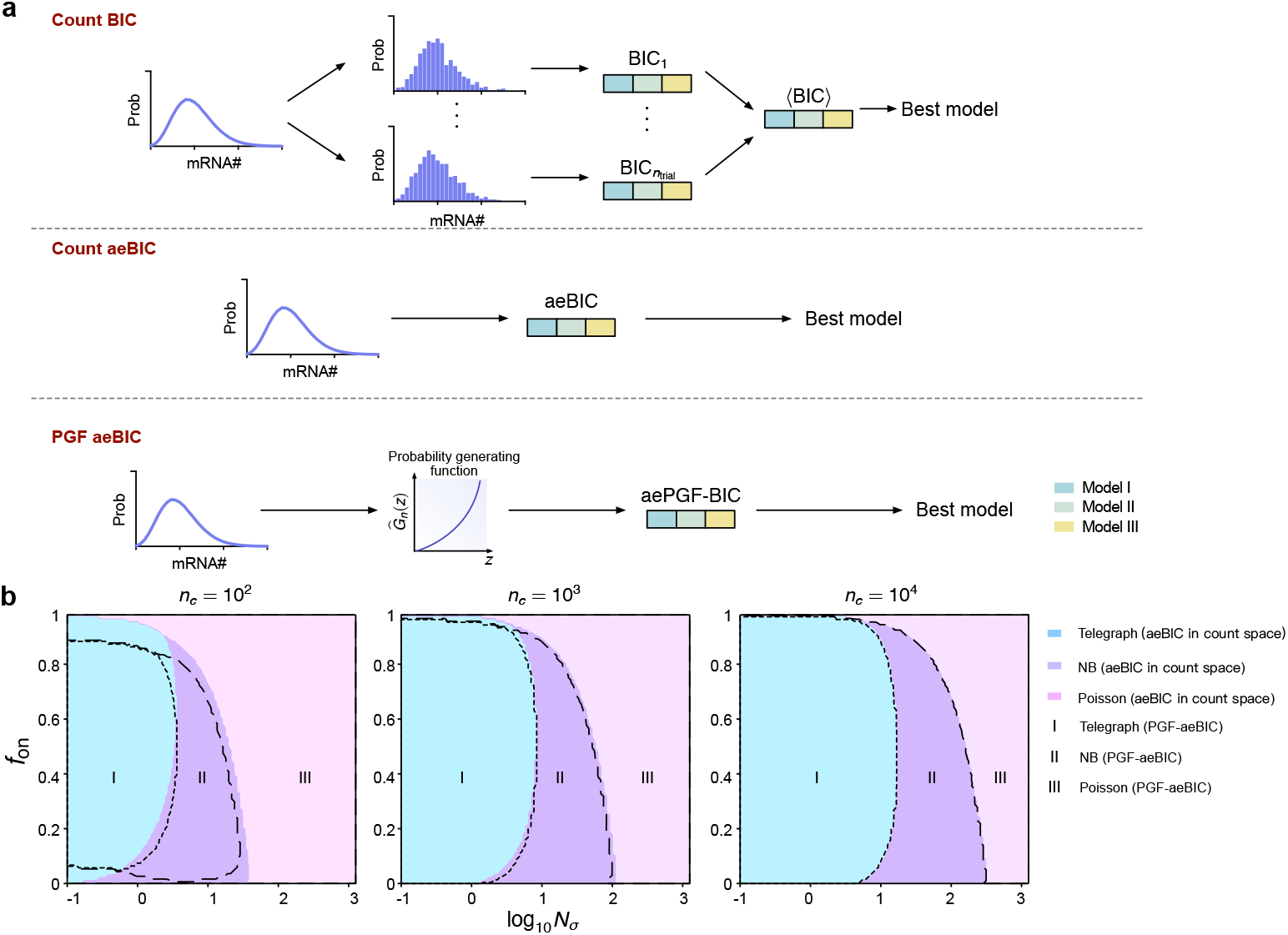
Comparison of count-space aeBIC and aePGF-BIC. (a) Schematic of the three model-selection procedures. Ordinary count-space BIC requires fitting each candidate to many independently sampled datasets and averaging the resulting scores to estimate ⟨BIC⟩. Countspace aeBIC replaces these repeated fits with a single population cross-entropy calculation. The proposed aePGF-BIC applies the same idea to the population PGF discrepancy. In every case, the candidate with the smallest score is selected. (b) Model-selection diagrams obtained by extending the design of Fig. 4 in Ref. [35] to PGF space. The stationary telegraph distribution is the generating distribution, the mRNA degradation rate is normalized to one, and the transcription rate is fixed at *ρ* = 15. The normalized switching speed *N*_*σ*_ = *σ*_on_ + *σ*_off_ and active-state fraction *f*_on_ = *σ*_on_*/N*_*σ*_ are varied, with results shown for *n* = 10^2^, 10^3^, and 10^4^ cells. At each grid point, the Poisson, NB, and telegraph candidates are fitted in count space by minimizing population cross-entropy and in PGF space by minimizing the population version of Eq. (2.4). All PGF candidates use the same collocation grid and covariance-weighting rule. The two sets of boundaries become more closely aligned as *n* increases, providing a population-level numerical illustration of the selection mechanism in Theorem 5.1.

Using the population log-likelihood *ℓ*_*r*_(***θ***_*r*_; ***ψ***) and the minimum KL discrepancy *κ*_*r*_(***ψ***) defined in Sec. 5, aeBIC can be written as

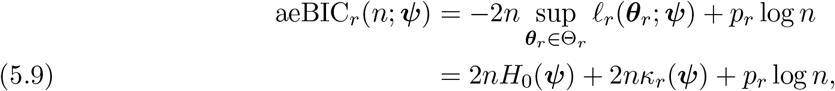

where ***ψ*** denotes the generating telegraph parameters and *H*_0_(***ψ***) is the entropy of the generating distribution. The entropy term is common to all candidates and therefore does not affect the selected model. The model-dependent part of aeBIC is consequently 2*nκ*_*r*_ + *p*_*r*_ log *n*, which is the population counterpart of the count-space expansion in Eq. (5.4). Because fitting each random sample can only improve its maximized likelihood, aeBIC is an upper population approximation to the BIC averaged over repeated samples. The remaining fitting advantage is lower order than the order-*n* discrepancy that separates a false model from the truth. See Fig. 3a for the workflow of aeBIC in count space.

The same construction can be carried out in PGF space. Recall from Eq. (4.2) that the PGF quasi-log-likelihood is a common normalization minus *nQ*_*r,n*_*/*2. Replacing the sample criterion by its population counterpart and minimizing over the candidate parameters gives the population PGF discrepancy δ_*r*_(***ψ***) in Eq. (5.2). Removing the normalization shared by all candidates then gives

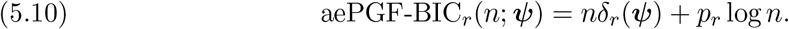

We call this quantity the approximate expected PGF-BIC (aePGF-BIC). With a fixed common weight, the expectation of the sample PGF criterion also contains the sampling-noise term tr{***W* Σ**_0_}. Because this term is the same for all candidates, it can be omitted without changing the selected model. When the weight is estimated from the data, Eq. (5.10) should be interpreted as the leading population approximation. The workflow of PGF-aeBIC is shown in Fig. 3a.

To compare the two population criteria, we extend the phase-diagram design of Fig. 4 in Ref. [35] to PGF space. The resulting phase diagrams in Fig. 3b show that aeBIC and aePGF-BIC can make different decisions at smaller sample sizes, especially near regions where two candidate distributions are difficult to distinguish. As *n* increases, the disagreement region becomes smaller, and the fraction of grid points assigned the same model by the two criteria increases. Thus, the count-space and PGF-space population criteria increasingly select the same Poisson, NB, or telegraph model as the sample size grows.

This result provides a partial numerical verification of Theorem 5.1. At a fixed regular interior parameter, the telegraph candidate has zero population discrepancy, whereas a separated Poisson or NB candidate has positive discrepancies in both count and PGF space. In aeBIC and aePGF-BIC, these discrepancies are multiplied by *n*, while the advantage of using fewer parameters grows only as log *n*. The lack-of-fit terms therefore eventually dominate the complexity advantage in both spaces, which is the same mechanism used in the proof of Theorem 5.1.

## 6. Cross-validation and its leave-one-out form in PGF space

The preceding sections selected models with information criteria. A second widely used model-selection method is cross-validation (CV), which asks how well each fitted candidate predicts observations that were not used to fit it. In *K*-fold CV, the cells are divided into *K* groups. Each candidate is fitted using *K* − 1 groups and evaluated on the remaining group. This is repeated until every group has served as validation data, the held-out errors are combined, and the candidate with the smallest validation error is selected. Leave-one-out cross-validation (LOO-CV) is the special case *K* = *n*. One cell is removed, each candidate is fitted to the other *n* − 1 cells, and its prediction for the omitted cell is evaluated. Repeating this operation once for every cell gives a model-selection score based entirely on held-out predictions. Although it requires *n* fits per candidate, each fit uses almost the full sample.

There is a classical connection between LOO-CV and information criteria in ordinary likelihood analysis (in count space). When held-out performance is measured by the log predictive density and each candidate is fitted by maximum likelihood, Stone showed that LOO-CV and the Akaike information criterion (AIC) give asymptotically equivalent model comparisons under regularity conditions and the specification condition used in that result [28, pp. 46–47, Eqs. (4.5)–(4.6)]. This is an AIC result, not a general equivalence between LOO-CV and BIC. AIC uses the complexity penalty 2*p*_*r*_, whereas BIC uses *p*_*r*_ log *n*. Nevertheless, LOO-CV and BIC can still choose the same model when that model has a fixed population fit advantage: the fit difference then accumulates in proportion to *n* and eventually dominates either penalty.

This raises the question relevant to the present framework: does the same large-sample agreement hold when both fitting and validation are performed in PGF space? Stone’s likelihood result does not answer this question directly, because Eq. (2.4) minimizes a covariance-weighted PGF discrepancy rather than a count-space negative log likelihood. We therefore construct a leave-one-out score from the same collocation grid, feature vector, and weighting rule as our PGF estimator, and then compare its selected model with PGF-BIC.

We now derive this PGF-space LOO score. Recall that cell *i* contributes the feature vector ***h***(*X*_*i*_). After removing that cell, the empirical PGF from the remaining cells is

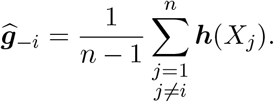

Let ***W***_−*i*_ be the model-independent weight calculated from those same training cells by the rule used for ***W***_*n*_. Candidate *r* is then fitted to the remaining cells by minimizing

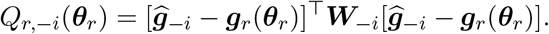

The resulting delete-one estimate is

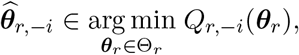

We next compare the omitted cell’s feature vector ***h***(*X*_*i*_) with the model PGF predicted from the other cells. Repeating this operation for every cell and adding the held-out errors gives the PGF-space LOO (PGF-LOO) score

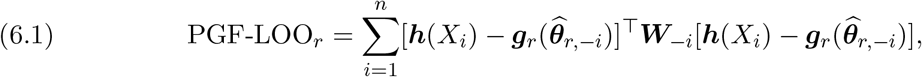

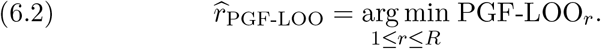

A smaller score means better prediction of omitted cells in PGF space. This is the usual idea of LOO prediction [28], applied to the same weighted PGF discrepancy used by Eq. (2.4) [10, 20]. No count-space likelihood is needed.

The next result gives one clear setting in which PGF-LOO and PGF-BIC eventually select the same model. Its central condition is that one candidate has a strictly smaller population PGF discrepancy than every competitor. In plain language, the best model must have a genuine fit advantage that does not shrink as more cells are collected.

### Theorem 6.1

(PGF-BIC and PGF-space LOO under finite-grid separation). *Assume that:*

1. *The cells are i*.*i*.*d. from P*_0_. *The candidate collection, collocation grid, and parameter dimensions p*_*r*_ *are fixed. Each parameter space* Θ_*r*_ *is compact, each model PGF* ***g***_*r*_ *is continuous, and every full-sample or delete-one fit is a measurable global minimizer of its criterion*.
2. *Removing one cell does not change the limiting weighting rule:*

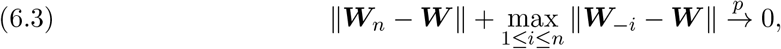

*where* ***W*** *is positive definite. This condition holds, for example, for a continuous regularized inverse of the cell-feature covariance when its population limit is nonsingular*.
3. *For every candidate r, the population criterion Q*_*r*_ *has one minimizer, denoted by* 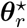. *Moreover, one candidate r*_⋆_ *has a strictly smaller optimized discrepancy than all the others:*

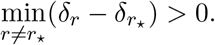

*Define*

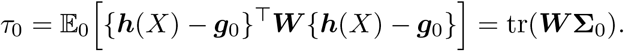

*This is the natural cell-to-cell variation in the PGF features. It remains present even when a model reproduces the population PGF exactly. Then*

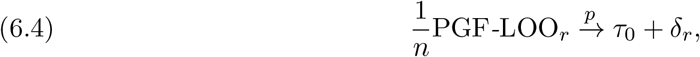

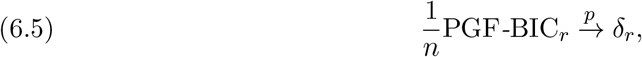

*simultaneously for all candidates in the finite collection. In particular*,

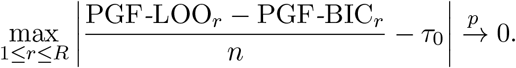

*The extra term τ*_0_ *changes the numerical value of PGF-LOO but is the same for every candidate because the weight is common across models. It therefore cannot change their ranking. Consequently*,

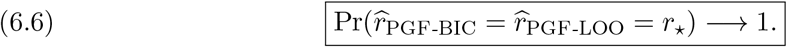

*A useful special case occurs when r*_⋆_ *is the only candidate that matches the population PGF on the chosen grid. The theorem also allows r*_⋆_ *to be the uniquely best approximation when every candidate is misspecified*.

*Proof*. The proof has three steps. First, omitting one cell has a vanishing effect on the empirical PGF. Second, all delete-one estimates approach the same population parameter. Third, we calculate the large-sample limits of the two scores.

We begin with the effect of deleting a cell. The following identity expresses the delete-one empirical PGF in terms of the full-sample empirical PGF:

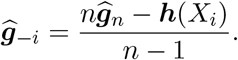

Every coordinate of ***h***(*X*_*i*_) lies in [0, 1], so 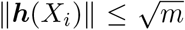. The same bound holds for the population PGF. It follows that

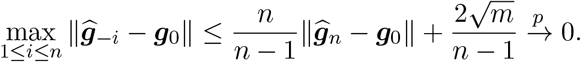

The first term tends to zero by the law of large numbers, and the second tends to zero directly. Thus deleting any one of the *n* cells has a vanishing effect, simultaneously for every possible omission. Denote the largest delete-one error by

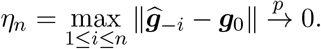

We next compare every delete-one criterion with the same population criterion *Q*_*r*_. Expanding the two quadratic forms and using 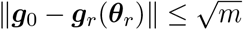 gives

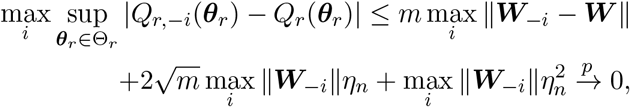

where Eq. (6.3) also gives max_*i*_ ∥***W***_−*i*_∥ = *O*_*p*_(1). Every term on the right tends to zero. Therefore, every delete-one sample criterion becomes uniformly close to *Q*_*r*_, for all parameter values and all omitted cells at once.

We now explain why the minimizing parameters must also be close. Fix any *ε >* 0. Compactness, continuity, and uniqueness imply that parameters at least *ε* away from 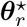 have a strictly larger population criterion. Write this positive gap as

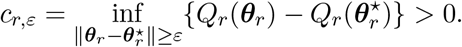

Suppose the uniform difference between *Q*_*r*,−*i*_ and *Q*_*r*_ is less than *c*_*r,ε*_*/*3. At 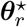, the delete-one criterion is at most 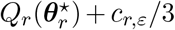. At a point outside the *ε*-ball, it is at least 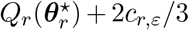. Every outside point therefore has a larger delete-one criterion and cannot be a minimizer. The conclusion holds for all omitted cells

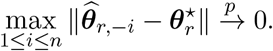

We next show carefully why the fitted model PGF in each held-out loss can be replaced by its population limit. Since ***g***_*r*_ is continuous on the compact set Θ_*r*_, it is uniformly continuous. Thus, for every *ε >* 0, there is a number *γ*_*ε*_ *>* 0, independent of *i*, such that

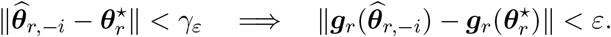

Because the same *γ*_*ε*_ works for every omitted cell, the preceding uniform convergence of the delete-one estimates implies

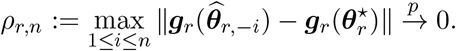

More explicitly, the event {*ρ*_*r,n*_ *> ε*} can occur only if at least one delete-one estimate is *γ*_*ε*_ or more away from 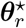. Therefore,

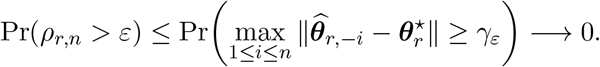

To translate this convergence into convergence of the held-out losses, define

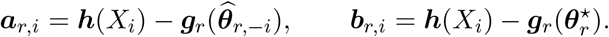

Every coordinate of ***h***(*X*_*i*_) and every coordinate of a PGF vector lies in [0, 1]. Hence

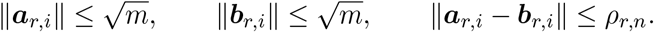

For each omitted cell *i*, let E_*r,i*_ denote the difference between the leave-one-out loss and its population-limit counterpart

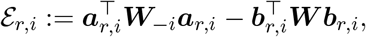

which can be expanded as

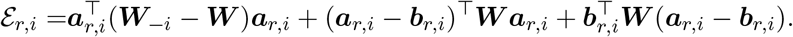

The first term changes only the weight matrix. The last two terms change the fitted PGF while holding the limiting weight fixed. Applying the standard matrix-norm inequality to these three terms gives

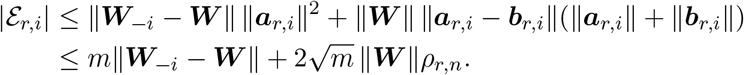

The first term tends to zero uniformly over *i* by Eq. (6.3), and the second tends to zero because 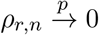. We have therefore proved, rather than merely assumed, that all held-out losses can be replaced at once:

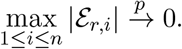

Taking an average cannot increase the largest approximation error, because

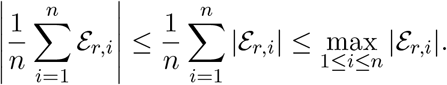

It follows that

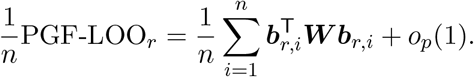

The expression remaining in the average now contains independent and identically distributed terms evaluated at the fixed population parameter 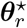. The law of large numbers therefore gives

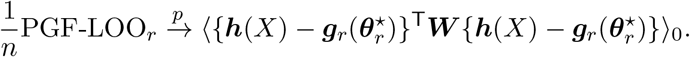

To evaluate this expectation, separate the held-out PGF error into two parts. Let

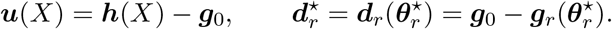

Here, ***u***(*X*) describes the random difference between one cell and the population PGF, whereas 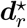 is the fixed PGF mismatch remaining after candidate *r* has been fitted as well as possible. Therefore,

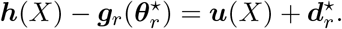

Substituting this decomposition into the expected held-out loss gives

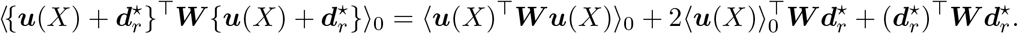

The first term measures natural cell-to-cell variation. Since ⟨***u***(*X*)⟩_0_ = **0** and ⟨***u***(*X*)***u***(*X*)^⊤^⟩_0_ = **Σ**_0_,

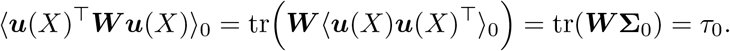

The cross term is zero because the random fluctuation is centered

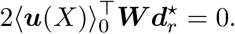

Finally, the last term is the smallest population PGF discrepancy achieved by candidate *r*:

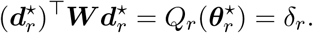

Combining the three terms gives

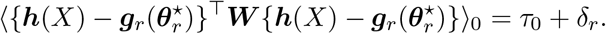

**Table 1:** Comparison of gene classifications from tenfold PGF cross-validation and PGF-BIC.

|  | Poisson (PGF-BIC) | Telegraph (PGF-BIC) | Total |
| --- | --- | --- | --- |
| Poisson (tenfold CV) | 769 | 748 | 1517 |
| Telegraph (tenfold CV) | 1199 | 4008 | 5207 |
| Total | 1968 | 4756 | 6724 |

This proves Eq. (6.4). The term *τ*_0_ is common to every candidate because it depends only on the true cell-to-cell variation and the common weight matrix. Only δ_*r*_ depends on the candidate model, so *τ*_0_ changes the numerical PGF-LOO scores but not their large-sample ranking.

There are only finitely many candidates, so the two limits hold for all of them simultaneously. PGF-LOO contains the additional *τ*_0_, but this quantity is identical for every candidate. Assumption 3 supplies a fixed positive gap between *r*_⋆_ and every competitor. With probability approaching one, the remaining sampling errors are too small to reverse that ranking. Both procedures therefore select *r*_⋆_, proving Eq. (6.6).

Theorem 6.1 establishes that PGF-BIC and PGF-space leave-one-out cross-validation select the same model under suitable large-sample conditions. Because tenfold cross-validation is more commonly used in practice, we compared it empirically with PGF-BIC using the MERFISH measurements of human osteosarcoma cells from Ref. [41]. The data were processed following p. 068401-9 of Ref. [36]. The original measurements contain nuclear and total mRNA counts for 10,050 genes across 1,368 cells. Low-expression genes and gene-specific outlier cells were removed, leaving 6,724 genes.

The data were processed following p. 068401-9 of Ref. [36]. The original measurements contain nuclear and total mRNA counts for 10,050 genes across 1,368 cells. Low-expression genes and gene-specific outlier cells were removed, leaving 6,724 genes. The candidates were a one-state constitutive model, denoted Poisson, and a two-state promoter-switching model, denoted Telegraph. Both models included the transportation of RNA from nucleus to cytoplasm and accounted for extrinsic variation by allowing transcription to depend on cell size. Because cell volumes were unavailable, the total transcript count in each cell was used as a proxy for cell size, following Ref. [36].

For PGF-BIC, both candidates were fitted to all available cells for each gene using the estimator in Eq. (2.4). The model with the smaller PGF-BIC value in Eq. (4.4) was selected. For tenfold cross-validation, the cells were divided into ten subsets. Each candidate was fitted to nine subsets and evaluated on the remaining subset, and this procedure was repeated for all ten folds. The candidate with the smaller median validation loss was selected.

The two procedures agreed for 4,777 of the 6,724 genes, giving an overall agreement rate of 71.0%. For the Telegraph classification, the 4,008 shared calls, together with 748 PGF-BIC-only and 1,199 cross-validation-only calls, give a precision of 0.843, a recall of 0.770, and an F1 score of 0.805. Thus, the two methods show strong concordance for Telegraph classifications, although agreement is weaker for Poisson classifications.

Unlike Ref. [36], we did not apply the one-standard-error rule [42] to the cross-validation results. That rule uses variation across folds to favor a simpler model when its performance is close to the minimum. PGF-BIC provides only one full-data score per candidate and has no directly comparable foldwise uncertainty without additional resampling. We therefore compared the unadjusted minimum PGF-BIC with the minimum median validation loss. This difference in selection rules partly explains why the classifications differ from those reported in Ref. [36]. These results are intended only as an empirical comparison of the two selection procedures. They should not be interpreted as evidence that a particular gene necessarily follows a unique biological mechanism.

## 7. Discussion

This work provides a unified way to perform parameter inference and model selection directly in PGF space. This is particularly useful when a model PGF is available analytically but computing the full stationary count distribution is difficult or expensive. The method also accounts for the fact that empirical PGF values evaluated at different points are calculated from the same cells and are therefore correlated. Most importantly, the same covariance-aware objective is used throughout: it estimates parameters and then supplies the fitted score used by PGF-BIC. Consequently, each candidate needs only one full-data fit, avoiding the repeated fitting required by cross-validation.

Theorem 3.1 explains why the parameter estimator is reliable. The empirical PGF is exactly centered on the true PGF. Under the stated regularity conditions, the estimated parameters approach their true values as more cells are observed, and their typical error decreases in proportion to *n*^−1*/*2^.

Theorem 5.1 shows that replacing the full count distribution with a fixed PGF summary need not change the final model decision. If the chosen PGF points distinguish every incorrect candidate, then PGF-BIC and ordinary count-space BIC select the same smallest correct model with increasing probability as the sample size grows. This concerns agreement of the selected model labels; the two criteria need not have the same numerical scores.

Theorem 6.1 establishes a related result for cross-validation: when one candidate has a uniquely smaller population discrepancy in PGF space, PGF-BIC and PGF-space leave-one-out cross-validation eventually select the same model. Consistent with this asymptotic result, the MERFISH analysis provides empirical evidence that PGF-BIC and tenfold PGF cross-validation can yield highly concordant model classifications in practice.

These results make PGF space a practical alternative to count-space analysis, but they do not remove the need for careful model design. The collocation grid must retain enough information to separate the candidates, and boundary or singular parameter regimes may require methods beyond ordinary BIC. Finally, a selected model should be interpreted as the simplest adequate description within the candidates considered, rather than as proof of a unique biological mechanism.

## Appendix A. Mathematical background for PGF-based inference

### Probability generating functions

A count distribution is usually described by the probabilities assigned to all possible counts. A probability generating function (PGF) packages the same information into the function

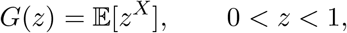

where *X* is a nonnegative integer-valued random variable. Knowing the full PGF determines the full count distribution. In practice, this paper evaluates the PGF at a finite collection of points *z*_1_, …, *z*_*m*_, called collocation points. These values form a compact summary of the count distribution and can often be calculated analytically even when evaluating every count probability is difficult. A finite grid does not necessarily distinguish every pair of distributions, so the choice of collocation points is part of the statistical model.

### Estimating the PGF from cells

For independent cell counts *X*_1_, …, *X*_*n*_, the PGF at *z* is estimated by averaging 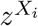 across cells:

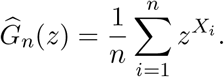

This empirical PGF is exactly unbiased: its expected value equals the population PGF for every sample size. Its sampling error decreases at the usual inverse-square-root rate as the number of cells grows.

Empirical PGF values at different collocation points are not independent, because all of them are calculated from the same cells. For example, a cell with an unusually large count influences every PGF coordinate simultaneously. The empirical PGF values must therefore be treated as one correlated vector. Their covariance matrix records both the uncertainty of each coordinate and the dependence between coordinates.

### Covariance-weighted parameter inference

For a large number of cells, the central limit theorem says that the empirical PGF vector is approximately multivariate Gaussian: its fluctuations have an approximately bell-shaped joint distribution whose covariance decreases as 1*/n*. This approximation leads to the Gaussian quasi-likelihood used in this paper. It is a likelihood approximation for the empirical PGF summary, not an exact likelihood for the original cell counts.

The estimator in Eq. (2.4) compares the empirical PGF vector with the vector predicted by a mechanistic model. Weighting this difference by the inverse covariance gives less influence to noisy directions and prevents strongly correlated coordinates from being counted as independent evidence. Under identification and smoothness conditions, the fitted parameters approach their population values, and their errors decrease in proportion to *n*^−1*/*2^. Exact finite-sample unbiasedness applies to the empirical PGF itself; the nonlinear parameter estimator is only first-order asymptotically unbiased under the additional tail condition stated in Theorem 3.1. The same construction extends to several molecular counts measured in each cell. The ordinary term *z*^*X*^ is then replaced by a product of powers, one for each count component, while the principles of cell averaging and covariance weighting remain unchanged.

### Model selection in PGF space

A more flexible model can generally reduce the fitted PGF discrepancy simply because it has more adjustable parameters. PGF-BIC corrects for this advantage by combining the optimized PGF discrepancy with the Bayesian information criterion (BIC) penalty *p*_*r*_ log *n*, where *p*_*r*_ is the number of identifiable parameters in candidate *r* and *n* is the number of independent cells. Thus, parameter inference and model selection use the same fit, and each candidate needs to be fitted only once.

The equivalence results in this paper concern selected model labels, not equality of numerical scores. When incorrect candidates remain separated from the truth on the chosen PGF grid, PGF-BIC and ordinary count-space BIC eventually select the same smallest correct candidate. Similarly, when one candidate has a uniquely smaller population PGF discrepancy, PGF-BIC and leave-one-out cross-validation performed in PGF space eventually choose that candidate. These conclusions need not hold when two nested models reproduce the chosen PGF values equally well. In that case, BIC increasingly favors the smaller model, whereas cross-validation uses a different complexity correction and may continue to select the larger model with positive probability.

A selected model should therefore be interpreted as the simplest adequate description among the candidates considered, relative to the chosen PGF grid and weighting rule. It is not, by itself, proof that the corresponding molecular mechanism is biologically unique.

## Data Availability

The MERFISH data analyzed in this study are publicly available in the supplementary data files `pnas.1912459116.sd12.csv` and `pnas.1912459116.sd14.csv` associated with Ref. [41].

## Artificial Intelligence Use

During the preparation of this work the authors used ChatGPT in order to improve the clarity and readability of the manuscript. After using this tool/service, the authors reviewed and edited the content as needed and take full responsibility for the content of the published article.

## Acknowledgment

This work is supported by NSFC Grants (62573195) and the Natural Science and Engineering Research Council of Canada’s (NSERC’s) Discovery Grant (RGPIN-2024-06015).

## Authorship and Contributorship Statement

All authors made substantial intellectual contributions to the conception, execution, and design of the work. All authors read and approved the final manuscript. The individual contributions were as follows:

- **Conceptualization:** Edward Z. Cao, Kim B. McAuley
- **Methodology:** Yiling Wang, Max Tomlinson.
- **Formal analysis and investigation:** Yiling Wang.
- **Software:** Yiling Wang, Zhanpeng Shu.
- **Data curation:** Zhanpeng Shu.
- **Visualization:** Yiling Wang, Zhanpeng Shu.
- **Writing—original draft preparation:** Yiling Wang, Edward Z. Cao.
- **Writing—review and editing:** Max Tomlinson, Zhanpeng Shu, Edward Z. Cao, Kim B. McAuley.
- **Funding acquisition:** Edward Z. Cao.
- **Resources:** Edward Z. Cao.
- **Supervision:** Edward Z. Cao.

